# The inner membrane complex protein, IMC55, is dispensable for intra-erythrocytic development of *Plasmodium falciparum*

**DOI:** 10.1101/2025.11.07.687246

**Authors:** Grace W. Vick, Manuel A. Fierro, Lenna Park, Carrie Brooks, Vasant Muralidharan

## Abstract

*Plasmodium falciparum*, an obligate intracellular Apicomplexan parasite and the causative agent of malaria must reside and replicate within a host cell during the intra-erythrocytic stage. The parasite utilizes a specialized double-membraned organelle, called the inner membrane complex for schizont-stage segmentation and merozoite motility during subsequent re-invasion. In this study, we investigate the function of IMC55, a multi-transmembrane inner membrane complex protein, which we identified as a putative interacting partner of the *P. falciparum* ER-resident calcium binding protein, PfERC. We have previously shown that PfERC is essential for parasite egress and our data shows that PfERC may also function in parasite invasion into the RBC. The whole genome transposon mutagenesis screen predicts that IMC55 is essential for the intra-erythrocytic asexual cycle. In this work, we show that IMC55 is not required for asexual replication. Following inducible expression of this gene using two conditional systems, we find there is no defect in replicative fitness during the asexual blood stage. Taken together, these data demonstrate that the function of IMC55 is either dispensable or redundant for the *Plasmodium falciparum* asexual blood stage.

## Introduction

Malaria is a deadly parasitic disease that kills around 600,000 people annually, with the highest burden of mortality in children under the age of five <u>(World Health Organization 2024)</u>. With over 263 million cases reported each year, this disease continues to pose a significant global health threat <u>(World Health Organization 2024)</u>. This life-threatening disease is caused by the eukaryotic parasite of the genus *Plasmodium.* Within this genus, *Plasmodium falciparum* overwhelmingly causes the most death, accounting for 95% of global malaria mortality <u>(Moxon et al. 2020)</u>. *P. falciparum* merozoites divide within red blood cells (RBCs) through a process called schizogony. This replicative cycle directly results in the manifestation of human clinical disease. To replicate, multiple rounds of asynchronous nuclear division precede segmentation and partitioning of subcellular contents to form nascent merozoites <u>(Francia and Striepen 2014)</u>. These daughter merozoites will egress and re-invade new host RBCs to perpetuate the cycle. Of these intra-erythrocytic infections, a small percentage of parasites will differentiate into either male or female form of the gametocyte <u>(K. A. Collins et al., 2018; Eichner et al., 2001)</u>. These transmissive-competent forms will be ingested by a mosquito during a blood meal and sexually reproduce to continue the infection cycle.

Throughout the entire life cycle, *P. falciparum* must undergo multiple dramatic changes in morphology and form in order to survive this wide range of host environments. To accomplish this feat, the parasite utilizes a highly specialized scaffold called the inner membrane complex (IMC) to orchestrate the need for structural integrity, cell division, and motility. This complex of flattened cisternal membrane compartments originates from the ancestral Alveolata lineage where it is shared among ciliates (termed alveoli), dinoflagellates (termed amphiesma), and Apicomplexans (termed IMC) <u>(Kono et al., 2013)</u>. This unifying membranous organelle functions in structural support for the parasite cell among all alveolates. In apicomplexans, the IMC has a broader function in segmentation during merozoite formation and anchoring the actin-myosin motor that powers gliding motility <u>(Keeley & Soldati, 2004; Kono et al.,</u>

2012).

The IMC is part of the three-bilayer membranous cytoskeleton, and consists of a series of flattened vesicles and interacting proteins that sit juxtaposed underneath the plasma membrane <u>(Morrissette & Sibley, 2002)</u>. These flattened sacs are termed alveoli and they are supported by a network of filamentous proteins that make up the subpellicular network (SPN) and underlying microtubules <u>(Mann & Beckers, 2001; Morrissette & Sibley, 2002)</u>. Together, the combined network of the parasite plasma membrane, IMC, and SPN form the pellicle that provides structural stability, anchoring of the glideosome, and scaffolding for daughter cells during segmentation <u>(Kono et al. 2013)</u>. The IMC is present in different forms through the various life stages as the parasite undergoes multiple metamorphoses, and its function and structure varies depending on the needs of each stage.

In this study, we investigate the function of an IMC protein (Pf3D7_0522600) across the asexual stage of the *P. falciparum* life cycle. Because this protein is localized within the IMC and has a molecular weight of 55 kDa, we’ve named it IMC55. We identified IMC55 through an interaction screen with the egress regulator endoplasmic reticulum (ER)-resident calcium binding protein (PfERC) <u>(Fierro et al. 2020; Kono et al. 2012)</u>. PfERC is a member of the CREC (calumenin, reticulocalbin 1 and 3, ERC-55, Cab-4) family of proteins, which have multiple EF hand domains and play various roles in calcium (Ca^2+^)-dependent processes in the secretory pathway <u>(Fierro et al., 2020; Honoré, 2009; Honoré & Vorum, 2000)</u>. We previously determined that PfERC functions in merozoite egress through the regulation of the aspartic protease, Plasmepsin X (PMX), which is essential for maturation of the subtilisin protease SUB1 <u>(Fierro et al., 2020)</u>.

IMC55 is a highly conserved, multi-pass transmembrane protein with ancient origins, sharing homologs across Apicomplexa and throughout Eukarya <u>(Kono et al. 2012)</u>. This protein has been localized to the IMC during asexual and sexual stages of the intra-erythrocytic development cycle, but no functional analyses have been performed. Previous phylogenetic analyses defined IMC55 to cluster in a non-alveolin, multi-transmembrane group of the inner membrane complex <u>(Kono et al. 2012)</u>. Observations of this protein during different stages of segmenting schizonts show that it has a specific localization pattern mirrored in glideosome-associated multi-membrane span proteins (GAPMs) <u>(Bullen et al. 2009; Kono et al. 2012; Hu et al. 2010)</u>. During early schizogony, this class of proteins form a cramp-like structure at the apical pole of the merozoite in a 2:1 ratio with the nucleus <u>(Hu et al. 2010)</u>. These structures then take on a ring-like form which becomes evenly distributed across the merozoite periphery <u>(Hu et al. 2010)</u>. This unique localization dynamic and the presence of multi-transmembrane domains might suggest a potential glideosome-associated function of IMC55 during schizogony. The genome-wide piggyBac mutagenesis screen in *P. falciparum* suggests that IMC55 is essential for the asexual life cycle <u>(Zhang et al. 2018)</u>. Similarly, transposon insertion mutagenesis predicts IMC55 as essential for the asexual life cycle of *P. knowesi* <u>(Elsworth et al. 2025; Oberstaller et al. 2025)</u>. However, a similar whole-genome essentiality screen in *P. berghei* found this gene to be dispensable <u>(Bushell et al. 2017)</u>. In this study, we generated conditional mutants of IMC55 to investigate its function during the asexual replicative stages within human RBCs.

## Results

### Identifying interacting proteins of PfERC

To identify interactors of PfERC, we utilized the previously established PfERC conditional mutant which was endogenously tagged with the inducible ribozyme, *glmS* (PfERC-*glmS*) or an inactive version of the ribozyme, *M9* (PfERC-*M9*) <u>(Fierro et al. 2020)</u>. Activation of the ribozyme through addition of glucosamine (GlcN) leads to cleavage of the active *glmS* from transcribed mRNA, resulting in target gene transcript degradation. The PfERC conditional mutants are tagged with a C-terminal 3X hemagglutinin (HA) tag, which was utilized to co-immunoprecipitate PfERC-associated proteins. Late schizonts were collected from PfERC-*M9* and untreated wild-type parental line (3D7) as a control, then lysed via sonication. PfERC interacting proteins were isolated from parasite lysates using anti-HA conjugated magnetic beads and identified via mass spectrometry analysis (Table S1). Abundance of each protein was measured and then averaged between biological replicates (Table S1). The ratio between protein abundance found in the PfERC IP compared to the control IP was calculated, and proteins enriched ≥ 10-fold in the PfERC IP were chosen for further study. There were 114 enriched proteins in the PfERC IP which were filtered for specific criteria (in blue, Fig. 1A). Since PfERC localizes to the parasite ER, we prioritized proteins with a predicted signal peptide and/or transmembrane domains <u>(Fierro et al. 2020)</u>. Candidates were further prioritized based on predicted essentiality, and peak expression during schizogony (Fig. 1B and 1C, Table 1) <u>(Zhang et al. 2018; Bushell et al. 2017)</u>. The final list is composed of 21 potential interacting partners of PfERC (Table 1).

**Figure 1.**
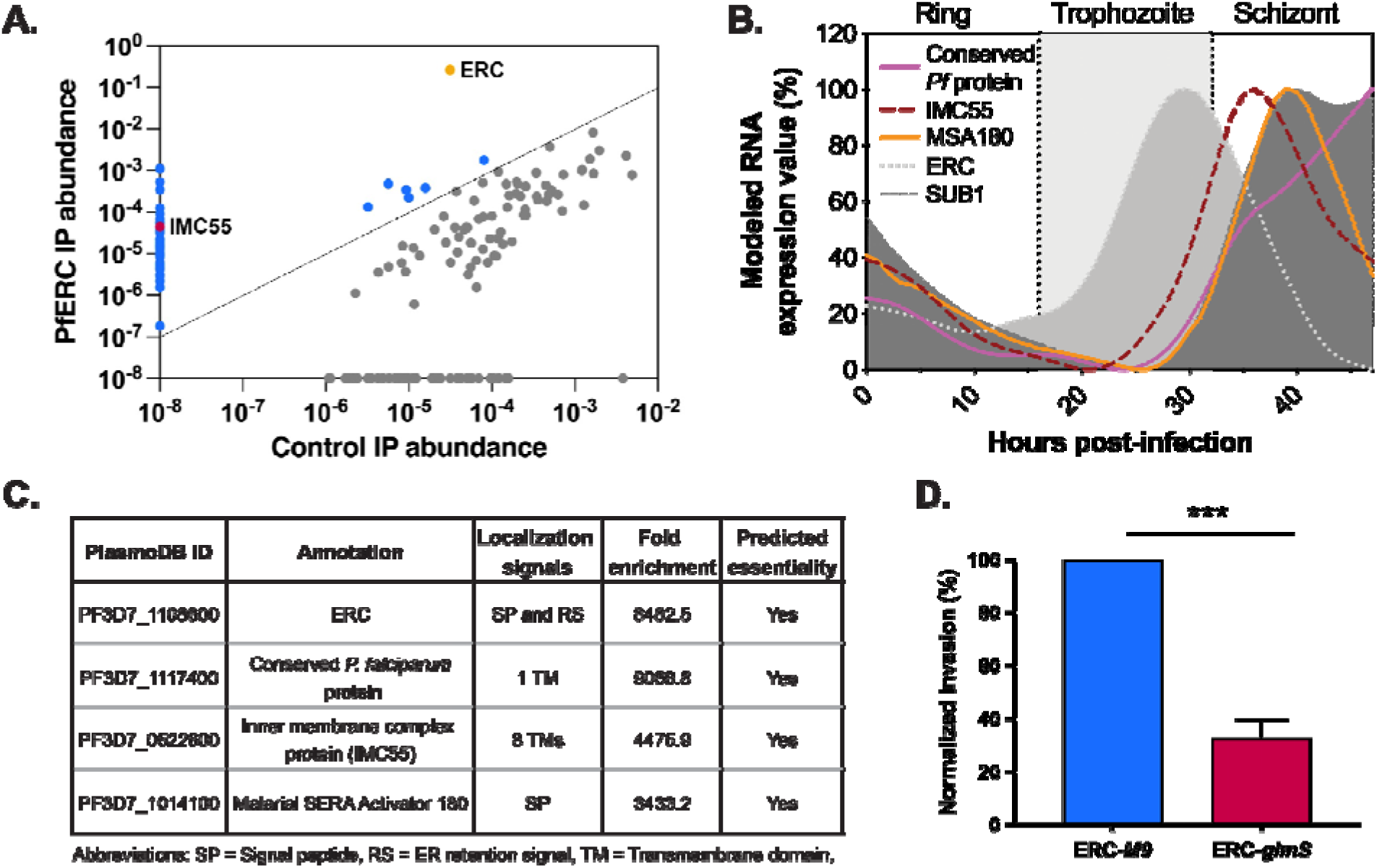
Identification of PfERC-interacting partners. (A) Plot of individual abundance values for all candidate proteins that passed specific criteria (blue) or 3D7 IPs (gray). PfERC is highlighted in yellow and inner membrane complex candidate protein IMC55 highlighted in pink. The threshold marked by the dotted line represents ≥ 10 fold enrichment in the PfERC IP. Data represented from 3 biological replicates, averaged together. (B) Graph showing mRNA expression during the 48 hour asexual development cycle of key PfERC-interacting candidates. (C) Fold enrichment and localization signals of PfERC-interactors of interest. (D) Quantification of PfERC-*glmS* and PfERC-*M9* invasion rate as shown in Supplemental Fig. 1. All samples were normalized to PfERC-*M9.* Error bars represent mean ± SEM. (n = 6 biological replicates; ***P<0.001, unpaired *t* test).

**Table 1.**
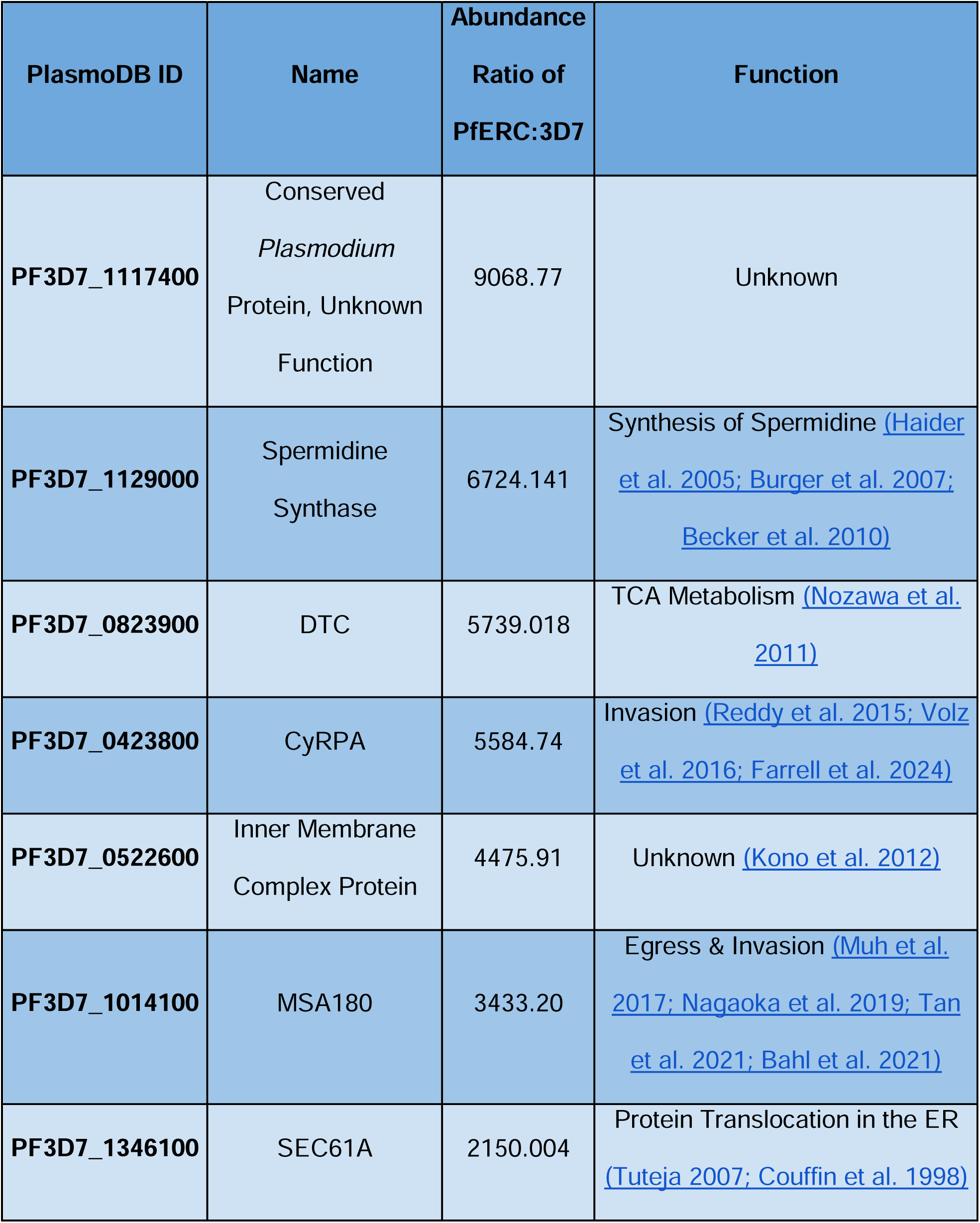
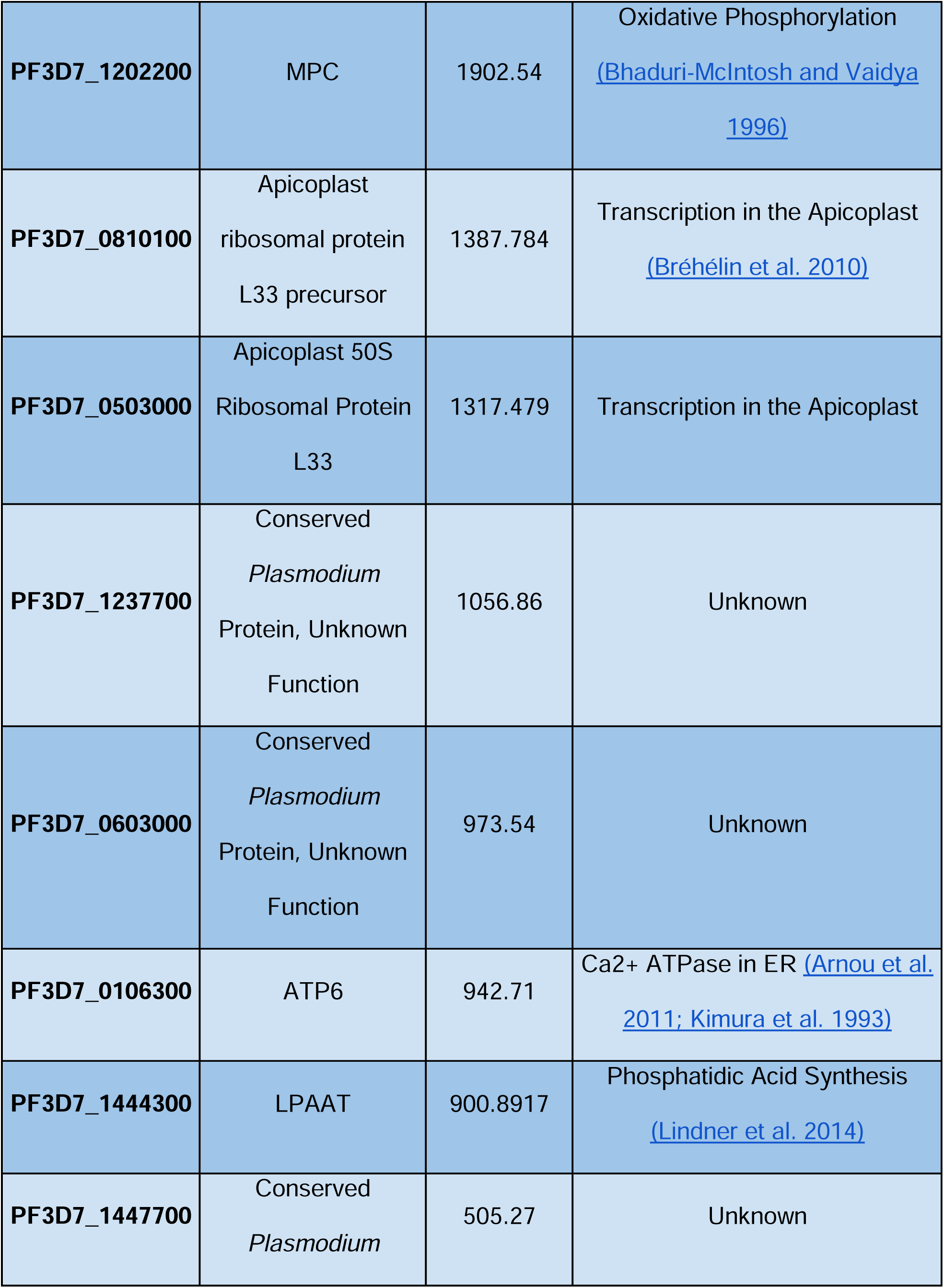
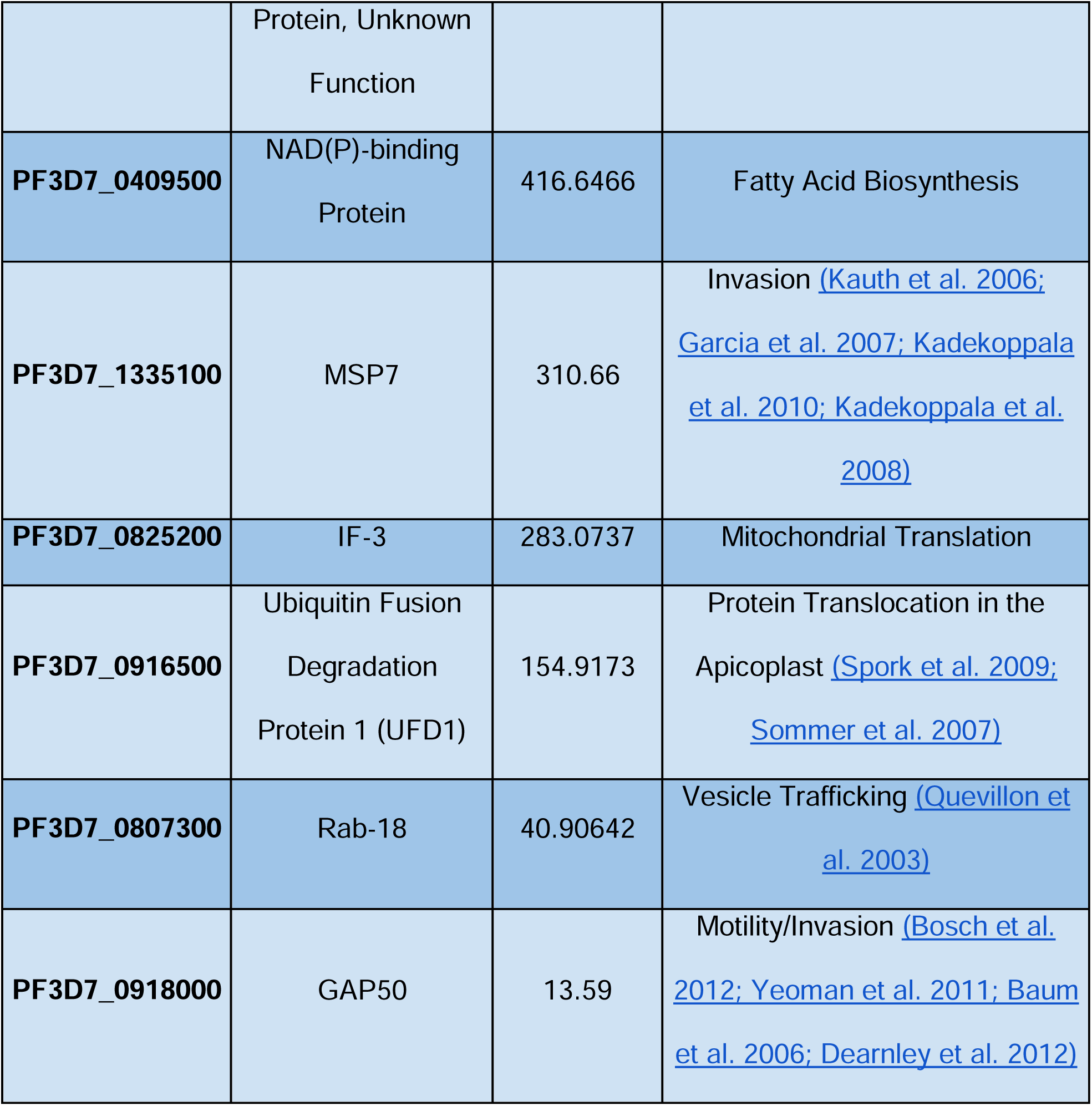
List of interacting partners of PfERC.

This list of PfERC-interactors includes proteins that were shown to function in merozoite egress, such as MSA180 (malarial SERA activator 180), which facilitates proteolytic processing of serine repeat antigen 6 (SERA6) that mediates RBCM rupture (Table 1) <u>(Bahl et al. 2021; Tan et al. 2021; Thomas et al. 2018; Ruecker et al. 2012)</u>. MSA180 has also been implicated as essential during invasion, as anti-MSA180 antibodies block invasion of merozoites, however this has not been corroborated <u>(Nagaoka et al. 2019)</u>. Interestingly, proteins that function in merozoite invasion, like cysteine-rich protective antigen (CyRPA) and merozoite surface protein 7 (MSP7) were identified in the PfERC IP (Table 1) <u>(Scally et al. 2022; Reddy et al. 2015; Farrell et al. 2024; Volz et al. 2016; Kadekoppala et al. 2008; Kauth et al. 2006)</u>. Additionally, this list includes two inner membrane complex proteins: the well-characterized glideosome-associated protein 50 (GAP50), and a protein with no known function (Pf3D7_0522600) (in pink, Fig. 1A and Table 1) <u>(Frénal et al. 2010; Perrin et al. 2018; Yeoman et al. 2011)</u>. Other notable proteins identified include ATP6 and spermidine synthase, in addition to ten other less-studied proteins, and four proteins with no known function (Table 1). Given the association between PfERC and several well-known invasion proteins as well as its requirement for PMX activity, we wanted to see if PfERC has a secondary function during invasion <u>(Fierro et al. 2020)</u>.

### PfERC is required for merozoite invasion

PfERC has previously been shown to regulate the egress of daughter merozoites through the maturation of PMX, an essential protease that processes crucial invasion ligands <u>(Fierro et al. 2020)</u>. Previous work had shown that PfERC-deficient parasites that managed to egress potentially failed to re-invade, as seen in Hema-3 stained thin-blood smears <u>(Fierro et al. 2020)</u>. To investigate this, we directly assessed the invasion efficiency of PfERC-deficient parasites. However, because PfERC has an essential role during egress, we required a technique that would isolate viable merozoites without the natural process of egress. To assess if PfERC has an additional downstream function during invasion, we used a mechanical syringe filter method to physically release infectious merozoites from egress-arrested schizonts <u>(Boyle et al. 2010)</u>. Taking synchronized PfERC-*glmS* and PfERC-*M9* late schizonts, we incubated parasites with 7.5 mM GlcN for 48 hours. Second cycle schizonts were stalled with the cysteine protease inhibitor E-64, which prevents RBCM rupture, trapping invasion-competent merozoites within the host cell <u>(Salmon 2001; Glushakova et al. 2009)</u>. After incubation with 20 μM E-64 for 8 hours, parasites were mechanically released through a syringe filter and allowed to invade fresh RBCs (Fig. S1A) <u>(Boyle et al. 2010)</u>. Using flow cytometry, we evaluated the invasion efficiency of PfERC-*glmS* parasites compared to the control PfERC-*M9* (Fig. S1A). We observe that PfERC-*glmS* parasites have a significantly reduced invasion rate with a ∼70% decrease in invasion efficiency, compared to the PfERC-*M9* parasites (Fig. 1D). These data show that PfERC has a secondary function during invasion, and when manually released from the RBC, merozoites lacking PfERC are unable to invade RBCs.

### IMC55 does not function in parasite replication

Given the requirement of PfERC for invasion, we were interested in discovering additional novel invasion regulators identified in the PfERC-interactors screen. We chose to study the function of IMC55 (Pf3D7_0522600) since the IMC is essential for successful invasion, serving as an anchor for the glideosome which powers motility during invasion (Table 1). To investigate the function of IMC55 during the intra-erythrocytic development cycle, we generated an inducible knockdown parasite line using the PfDOZI-*TetR* aptamer system (termed IMC55^apt^) (Fig. 2A) <u>(Rajaram et al. 2020; Ganesan et al. 2016)</u>. We utilized CRISPR/Cas-9 engineering to modify the endogenous target locus, incorporating into the repair template a Blastacidin-S (BSD) drug selection cassette, a 3X HA epitope tag, and the 10X *tetR* aptamer (Fig. 2A). Integration of the modified gene was confirmed via PCR using integration-specific primers (Fig. 2B and Table S2). To confirm subcellular localization of the IMC55^apt^ mutant, we used immunofluorescence assay (IFA) and co-stained fully segmented schizonts with anti-HA and a marker of the merozoite membrane surface, merozoite surface protein 1 (MSP1). As the IMC sits within ∼20-30 nm below the parasite plasma membrane, few tools have the resolution to differentiate these membranes <u>(Clements et al. 2022)</u>. Previous studies have shown MSP1 to localize just above the IMC making it a strong proxy for an IMC marker <u>(Liffner et al. 2023)</u>. We observe that IMC55 outlines each segmented merozoite just near the parasite plasma membrane, agreeing with previous reports of localization to the IMC (Fig. 2E) <u>(Kono et al. 2012)</u>.

**Figure 2.**
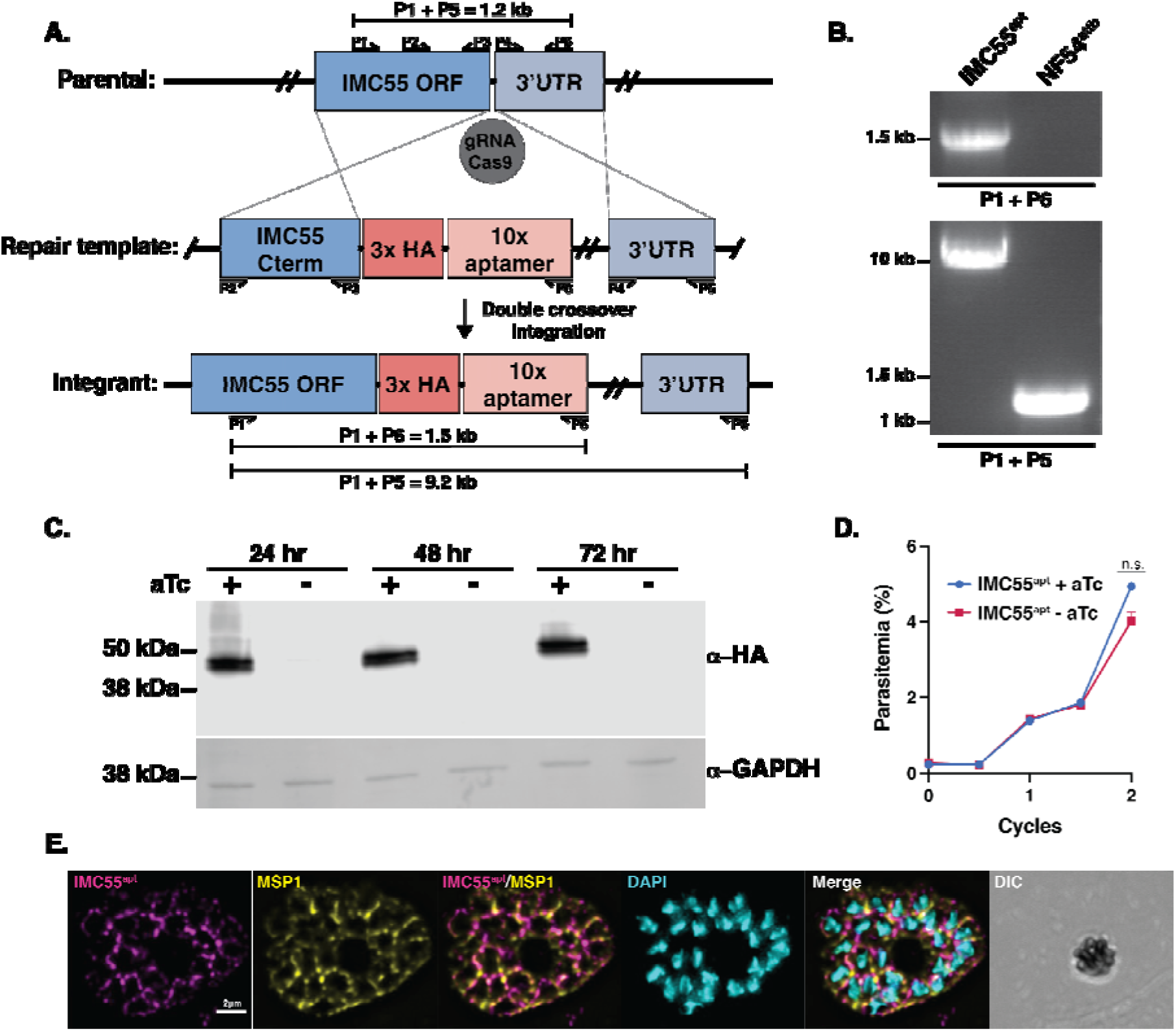
Generation of IMC55^apt^. (A) Schematic showing integration of the repair plasmid used to incorporate the *TetR* aptamer system into the endogenous IMC55 genomic locus. (B) PCR amplification showing integration of modified IMC55^apt^ locus. Primer pairs are specific for amplifying the region between the C-terminus and 3’UTR or between the C-terminus and integrated 10X aptamer repeats. (C) Western blot of IMC55^apt^ lysates after 24, 48, or 72 hours grown with or without 500 nM aTc. The membrane was probed with anti-HA to detect IMC55^apt^ protein, or anti-GAPDH as a loading control. (n = 3 biological replicates). (D) Growth assay of IMC55^apt^ parasites in the presence or absence of 500 nM aTc for two complete intra-erythrocytic life cycles, measured via flow cytometry. Representative of 3 biological replicates. Each data point represents the mean of three technical repeats. (Error bars = SD; not significant by 2-way ANOVA). (E) Immunofluorescence assay showing subcellular localization of IMC55^apt^. Antibodies used were anti-HA to detect IMC55^apt^, a marker for the merozoite surface (anti-MSP1), and a DNA stain (DAPI). Images include merged fluorescence and phase contrast (DIC). Images were collected as a Z-stack, deconvolved, then shown as a maximum intensity projection. Representative images from 3 biological replicates.

With the inducible *tetR* aptamer system, expression of IMC55 is dependent on the presence of anhydrotetracycline (aTc), a tetracycline analog <u>(Rajaram et al. 2020)</u>. To confirm that expression of IMC55 requires aTc, we cultured tightly synchronized IMC55^apt^ ring-stage parasites with or without 500 nM aTc and collected lysates at 24, 48, and 72 hours. We evaluated protein expression of these lysates via western blot and observed over 97% reduction in IMC55^apt^ expression by 24 hours post-aTc removal (Fig. 2C).

Next, we evaluated if IMC55 localization relies on signal-dependent exocytosis of egress-specific vesicles. It’s been shown that upstream PKG signaling is responsible for exocytosis of the egress-specific vesicles, exonemes, and the invasion-specific vesicles, micronemes <u>(Brochet et al., 2014; Dedkhad et al., 2025; Dvorin et al., 2010; Yeoh et al., 2007)</u>. This proteolytic cascade is responsible for translocation of specific proteins such the invasion ligand, apical membrane antigen 1 (AMA1) <u>(Absalon et al., 2018; C. R. Collins, Hackett, et al., 2013)</u>. To assess if translocation of IMC55 requires PKG, we inhibited PKG signaling using the specific PKG inhibitor compound 1 (C1) which stalls the proteolytic cascade, blocking downstream exoneme exocytosis and egress <u>(Collins et al. 2013; Dedkhad et al. 2025)</u>. We took synchronized IMC55^apt^ schizonts at 46 hours post invasion and incubated them with 1.5 µm C1. After a four-hour incubation, we collected samples for IFA. When PKG signaling is inhibited, we observed that IMC55^apt^ has the same localization pattern as untreated conditions, indicating that IMC55 is constitutively present in the IMC and does not translocate prior to egress (Fig. S2A).

Given the strong efficacy of this knockdown system (Fig. 2C), and the suggested essentiality of IMC55 in the whole-genome transposon screen, we wanted to re-evaluate the essentiality of IMC55 for intra-erythrocytic development <u>(Zhang et al. 2018)</u>. To assess the consequence of IMC55 knockdown on asexual replication, we measured growth of parasites over two complete intra-erythrocytic replicative cycles using a flow cytometry-based growth assay. We took synchronized IMC55^apt^ ring-stage parasites and incubated them with or without 500 nM aTc for four days, measuring parasitemia every 24 hours. Surprisingly, we found no difference in the asexual expansion of IMC55^apt^ grown with or without aTc (Fig. 2D). Together, these data show that tagging of IMC55 does not alter its subcellular localization, and the conditional knockdown of IMC55 expression does not result in a growth defect during the asexual blood stages (Fig. 2).

### Conditional knockout of IMC55 does not inhibit asexual growth

The transposon insertional mutagenesis data suggest that IMC55 is essential for the asexual life cycle, but the experimental data using the conditional knockdown system did not show any replicative fitness defect <u>(Zhang et al. 2018)</u>. Therefore, to confirm if IMC55 is essential for the asexual blood stages, we generated an inducible knockout (KO) of IMC55 (IMC55^KO^) by flanking the coding region with *loxp* (LoxPint) sites and transfecting into the NF54::DiCre parental line that expresses both halves of the rapamycin-dimerizing Cre recombinase (Fig. 3A) <u>(Collins et al. 2013; Kudyba et al. 2021; Jones et al. 2016)</u>. The major advantage of utilizing this system is that the entire gene can be excised, completely preventing transcription in a highly efficient manner. Using CRISPR/Cas-9, we modified the endogenous locus to contain a hDHFR drug selection cassette and a 3X HA epitope tag (Fig. 3A). Integration of the DiCre system was confirmed via PCR using integration-specific primers (Fig. 3B and Table S2). To verify that excision of IMC55 is dependent on rapamycin addition, we cultured tightly synchronized IMC55^KO^ ring-stage parasites with 100 nM rapamycin (RAP) or 0.05% dimethyl sulfoxide (DMSO) for 4, 8, or 16 hours and collected lysates for PCR amplification immediately post-RAP treatment. We observe that 4-hour and 8-hour incubations with RAP are not sufficient to completely excise the gene, as faint bands corresponding to the un-excised amplicon are present (Fig. S3A). However, a 16-hour incubation with RAP is adequate to completely excise the gene (Fig. S3A). After a 16-hour RAP-treatment, excision of the floxed IMC55^KO^ results in amplification of a smaller band size (3.1kb) compared to the DMSO-treated IMC55^KO^ (7.8kb) or parental NF54::DiCre line (3.8kb) (Fig. 3A and 3B). These data show that IMC55 is efficiently excised after rapamycin addition (Fig. 3B).

**Figure 3.**
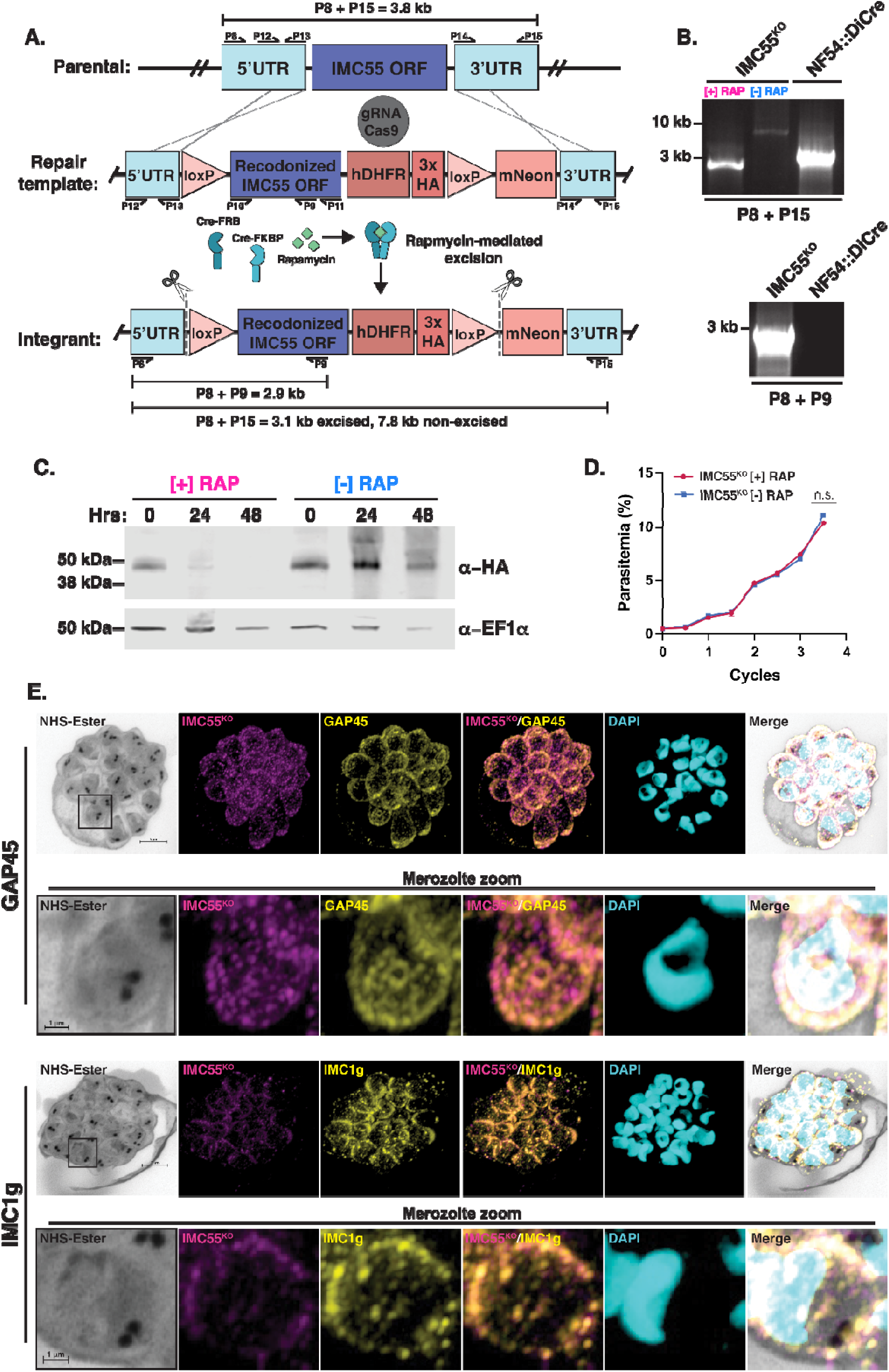
IMC55^KO^ is not essential for intra-erythrocytic development. (A) Schematic showing the repair plasmid template to introduce the DiCre knockout system into the IMC55 genomic locus. (B) PCR test showing integration of modified IMC55^KO^ locus. Primer pair P8 + P15 are specific to amplifying the region between the 5’UTR and 3’UTR. In the presence of rapamycin, the excised amplicon is 3.1kb, and in the absence of rapamycin, the amplicon is 7.8kb. The parental unmodified amplicon is 3.8kb. Primer pair P8 + P9 amplifies the region between 5’UTR and recodonized IMC55 ORF. (C) Western blot showing IMC55^KO^ expression after treatment with 100 nM rapamycin or 0.05% DMSO for 16 hours, with lysates collected at 0, 24, or 48 hours. Membrane was probed with antibodies against HA to detect IMC55^KO^, or anti-EF1α (loading control). (n=3 biological replicates). (D) Growth assay of IMC55^KO^ after treatment with 100 nM rapamycin or 0.05% DMSO for 16 hours, measured via flow cytometry. Representative of 4 biological replicates. Each data point represents the mean of three technical repeats. (error bars = SD; not significant by 2-way ANOVA). (E) Ultrastructural expansion microscopy (U-ExM) of IMC55^KO^ colocalization with inner membrane markers. ML10-arrested late schizonts were expanded by U-ExM, fixed, then stained with anti-HA (magenta), anti-GAP45 (yellow, top), anti-IMC1g (yellow, bottom), NHS-Ester (grayscale), and the DNA stain SYTOX (cyan). Representative images were collected as a Z-stack, deconvolved, then selected slices were projected as a single image. Scale bar =1 μm.

IMC55 has been shown to co-localize with the well-characterized protein GAP45 during schizogony, an integral glideosome protein that bridges the gap between IMC and parasite plasma membrane <u>(Frénal et al. 2010; Baum et al. 2006; Jones et al. 2009; Rees-Channer et al. 2006; Green et al. 2008; Kono et al. 2012)</u>. Therefore, we stained fully segmented IMC55^KO^ schizonts with the IMC marker GAP45 (Fig. S4) <u>(Baum et al. 2006; Jones et al. 2006; Kono et al. 2012; Ridzuan et al. 2012)</u>. IMC55^KO^ colocalizes with GAP45 as expected, corroborating previously observed reports (Fig. S4A) <u>(Kono et al., 2012)</u>.

Given the limiting resolution of conventional immunofluorescence microscopy, we used ultrastructural expansion microscopy (U-ExM), to reveal greater subcellular detail of IMC55 localization. U-ExM is a method that physically expands the sample ∼4.5X, which provides increased clarity of subcellular organelle morphology <u>(Liffner and Absalon 2021)</u>. To capture fully mature, segmented schizonts, the PKG inhibitor ML10 which stalls schizont egress was applied <u>(Ressurreição et al., 2020)</u>. Expanded ML10-arrested IMC55^KO^ schizonts were stained with the IMC markers anti-GAP45 and anti-IMC1g, in addition to the amine-reactive NHS-Ester stain. NHS-Ester allows us to visualize protein-rich cellular content such as the rhoptries, basal complex, and apical polar rings. In these images, IMC55 strongly patterns as ring-like structures completely surrounding the daughter merozoites. This localization mirrors the patterning observed for both IMC markers in fully segmented schizonts (Fig. 3E). The patterning of IMC55 seems less distinct, with a more diffuse and slightly punctate expression around each merozoite compared to either IMC marker (Fig. 3E, merozoite zoom). The localization of IMC55 observed via U-ExM agrees with our previous data (Fig. 2E and S4A) and prior observations <u>(Kono et al. 2012)</u>.

To evaluate the conditional expression of protein, tightly synchronized IMC55^KO^ ring-stage parasites were treated with or without 100 nM rapamycin and collected for saponin lysis at 0, 48, and 96 hours. IMC55^KO^ protein expression was quantified using western blot and was found to have > 92% reduction in RAP-treated IMC55^KO^ expression at 48 hours and almost 99% reduction by 96 hours post treatment (Fig. 3C). With this robust conditional knockout system, we next wanted to interrogate if IMC55 is truly dispensable for asexual replicative fitness. Synchronized ring-stage IMC55^KO^ parasites were incubated with 100 nM rapamycin or 0.05% DMSO for 16 hours and evaluated every 24 hours for seven days (3.5 asexual life cycles). Using a flow cytometry-based growth assay, we measured parasitemia over the multiple replicative cycles. Interestingly, RAP-treated IMC55^KO^ parasites had no significant difference in parasitemia compared to DMSO-treated IMC55^KO^ (Fig. 3D). This observation agrees with our earlier findings of IMC55^apt^ knockdown. Together, these data establishes IMC55 as a non-essential member of the IMC during the asexual replicative cycle.

## Discussion

During asexual development, the IMC plays a key role in the genesis of progeny and motility of invasive forms <u>(Ferreira et al. 2020)</u>. Moreover, this membranous organelle provides the structural support necessary for multiple parasite remodeling events across the life cycle <u>(Harding and Meissner 2014)</u>. Although our knowledge of this unique structural organelle has increased over time, how the IMC functions to support the varying needs of the parasite is still an area of poor understanding <u>(Ferreira et al. 2020)</u>. Determining which IMC-localized proteins are indispensable or redundant is key in advancing our knowledge of IMC biology across the entire life cycle.

Previous work has identified the essential ER-resident calcium-binding protein PfERC that is responsible for the proteolytic processing of SUB1 and PMX during egress. To identify PfERC-interacting proteins, we performed a co-immunoprecipitation screen which unveiled associations with both well-characterized and novel proteins essential for egress, invasion, and protein translocation (Table 1). In addition to the work presented here, this PfERC-interactor candidate list may uncover previously unknown genes which are essential effectors of the asexual blood stage. Interestingly, PfERC seems to associate with a number of known invasion effectors such as CyRPA, MSP7, and GAP50 (Table 1). CyRPA is an essential component of the pentameric PCRCR complex that forms a bridge between the invading merozoite and erythrocyte membrane <u>(Scally et al. 2022; Reddy et al. 2015; Farrell et al. 2024; Volz et al. 2016)</u>. MSP7, a surface protein found in complex with MSP1 and MSP6, has a minimal, yet functional role in invasion <u>(Kauth et al. 2006; Garcia et al. 2007; Kadekoppala et al. 2010; Kadekoppala et al. 2008)</u>. GAP50 is an integral IMC protein that secures the glideosome machinery into the IMC membrane <u>(Johnson et al. 2007; Gaskins et al. 2004; Yeoman et al. 2011)</u>. Taken together, these interactions suggest that PfERC might have an additional specific function in the invasion pathway.

These data led us to investigate if PfERC has a secondary function during invasion that was not observed due to its upstream role regulating egress <u>(Fierro et al. 2020)</u>. We found that PfERC does have an essential function during invasion, and when mechanically freed from host cells, PfERC-deficient merozoites were not invasion-competent. These data corroborate previous work showing that PfERC regulates proteolytic processing of the invasion ligands MSP1 and AMA1 <u>(Fierro et al. 2020)</u>. Given that these invasion proteins are processed prior to egress, the invasion defect seen in the PfERC knockdown is likely a secondary effect <u>(Nasamu et al. 2017; Pino et al. 2017; Das et al. 2015; Thomas et al. 2018)</u>. In light of PfERCs dual function in egress and invasion, elucidating its downstream role during invasion will be technically challenging. With the current limitations, we would need to utilize a specific, PfERC-targeted small molecule such as an endoperoxide to disentangle the distinct role of PfERC during invasion <u>(Morita et al. 2012)</u>.

Interestingly, an IMC protein, IMC55, was revealed as a potential interactor of PfERC (Table 1). Given the identified association between these proteins but disparate localization, it’s likely that the interaction found in the screen is transient while IMC55 is being transported through the ER to its final destination. Previous studies have described IMC55 as a novel IMC protein, but its biological function remains unknown <u>(Kono et al. 2012)</u>. In this work, we utilized CRISPR/Cas9 gene editing to elucidate the functional mechanism of IMC55. First, we chose to use the *TetR* aptamer conditional knockdown system that allows us to control gene expression in an inducible and reversible manner (Fig. 2A) <u>(Rajaram et al. 2020)</u>. Following the removal of aTc, there was a strong reduction in protein expression indicating the knockdown system was highly efficient (Fig. 2C). However, when measuring the replicative growth of IMC55-deficient parasites, we found there was no replicative defect compared to the control.

Since conditional knockdown of IMC55 did not inhibit parasite growth, it’s likely that either the transposon-based screen resulted in a false positive in the case of IMC55 or that the residual expression in the IMC55^apt^ (∼3% in the absence of aTc) was sufficient for parasite survival (Fig. 2D). This has been observed before for other essential proteins such as plasmepsin V (PMV) and caseinolytic protease (ClpP), where residual expression upon knockdown led to no growth defects and using more robust conditional knockout and knockdown (>99% knockdown) showed that these proteins were indeed essential for parasite survival <u>(Florentin et al. 2017; Sleebs et al. 2014; Boonyalai et al. 2018; Florentin et al. 2020; Polino et al. 2020)</u>. To verify that IMC55 is dispensable for the asexual blood stage, we employed the conditional knockout DiCre system which relies on the addition of rapamycin for full excision of the target gene <u>(Jones et al. 2016)</u>. Complete excision of the IMC55 gene required a 16-hour incubation with rapamycin, instead of the standard 4-hour incubation as previously described <u>(Jones et al. 2016)</u>. However, within 48 hours of rapamycin treatment, we observe > 99% reduction in expression of IMC55^KO^, indicating this system is incredibly robust (Fig. 3C). To thoroughly evaluate replication in IMC55-depleted parasites, the growth assay duration was extended in RAP-treated IMC55^KO^ (3.5 cycles), compared to the conditional knockdown (2 cycles) (Fig. 2D and 3D). Even with this highly robust knockout system, we demonstrate that IMC55 is dispensable for the asexual blood stage. Together, the current study uses multiple strategies to establish that IMC55 has a non-essential or redundant role in the IMC during intra-erythrocytic asexual replication.

In *P. falciparum*, a genome-wide piggyBac mutagenesis screen predicted IMC55 to be essential, but the screen was not completed to saturation and no transposons were inserted into the IMC55 coding sequence <u>(Zhang et al., 2018)</u>. It’s likely that the lack of transposon insertions into IMC55 resulted in a false-positive mutagenesis index score <u>(Zhang et al. 2018)</u>. While gene disruption methods such as *Plasmodium* genetic modification (PlasmoGEM) and piggyBac transposon mutagenesis are powerful genetic screening tools, they are not infallible to false-positive and false-negative artifacts <u>(Martin 2020)</u>. Together, we have experimentally shown that IMC55 is either a truly non-essential component of the inner membrane complex or more likely, is functionally redundant during the asexual blood stage. Currently, the data presented here cannot distinguish between these possibilities, and with no observable phenotype following IMC55 knockout, the function of this gene is challenging to determine. However, while not assessed in this study, IMC55 may be essential for a different stage of the *P. falciparum* life cycle such as in the mosquito or liver. Interestingly, the Rodent Malaria gene modification screen performed in *P. berghei* predicted PbIMC55 (PBANKA_1237300) to be dispensable for gametocyte development <u>(Bushell et al. 2017)</u>. Further studies would need to be performed to determine if IMC55 is required for other stages.

IMC55 is a highly conserved protein within Eukarya, with orthologs across Alveolates, Fungi, and Viridiplantae <u>(Kono et al. 2012)</u>. Within *Plasmodium* IMC proteins, this high level of conservation is not universally found <u>(Kono et al. 2012)</u>. Some IMC members such as IMC55 and IMC1g share ancient origins, while other integral proteins such as GAPM3 and IMC1j are only found within Apicomplexa <u>(Kono et al. 2012)</u>. While we were unable to determine the functional role of IMC55, the InterPro PFAM database predicts this gene to contain a magnesium (Mg^2+^) transporter (NIPA) domain <u>(Paysan-Lafosse et al. 2025)</u>. NIPA (nonimprinted in Prader-Willi/Agelman syndrome) is a family of transmembrane proteins that selectively mediate Mg^2+^ ion transport <u>(Goytain et al. 2007; Goytain et al. 2008)</u>. Magnesium is a prevalent cation that is a critical co-factor in many enzymatic and physiological processes such as protein and nucleic acid synthesis, energy production, and membrane stabilization <u>(Franken et al. 2022)</u>. In *Plasmodium*, most Mg^2+^ transporters localize within the mitochondria, and few have been functionally analyzed <u>(Wunderlich 2022)</u>. Of the ∼50 IMC proteins identified, the only protein with a predicted Mg^2+^ transporter functional domain is IMC55 <u>(Ferreira et al. 2020; Paysan-Lafosse et al. 2025; Wichers et al. 2021)</u>. In a Turbo-ID-based proteomic study in *P. yoelii*, over 300 candidate IMC proteins were identified, of which the only *P. falciparum* ortholog with putative Mg^2+^ transport function was IMC55 <u>(Qian et al. 2022)</u>. Given the lack of phenotype when disrupted, we were unable to evaluate the functional role of IMC55 as a putative Mg^2+^ transporter. However, if IMC55 has a functionally redundant role in Mg^2+^ transport, no other IMC proteins have yet been identified that might compensate in this biochemical pathway during IMC55 knockout.

Given the widely conserved nature of this protein, it is curious that the data show that IMC55 is dispensable for the *Plasmodium* intra-erythrocytic development cycle. Functional analyses across IMC55 orthologs are sparse, even in related apicomplexan *Toxoplasma gondii*, where the IMC55 ortholog (TGME49_312622) is predicted to be essential for the lytic cycle <u>(Sidik et al. 2016)</u>. In a BioID-based study to identify novel IMC proteins in *T. gondii*, TgIMC55 was identified as an interactor of the IMC protein ISP3 <u>(Chen et al. 2015)</u>. However, no experimental analyses have been performed to elucidate the molecular function of TgIMC55.

In summary, our work has built on the previous identification of the novel inner membrane complex protein, IMC55. To evaluate the functional role of IMC55 during asexual replication, we chose to employ multiple genetic editing strategies. Together, our data shows that IMC55 is dispensable or functionally redundant during the intra-erythrocytic asexual replicative stages.

## Methods and materials

### Generation of plasmids

Genomic DNA was isolated from NF54^attB^ parasites using the QIAgen DNeasy Tissue & Blood extraction kit. Primer pairs P2 & P3 were designed to amplify a ∼450 bp fragment of the IMC55 C-terminus (Pf3D7_0522600), and primer pairs P4 & P5 were used to amplify a ∼450 bp fragment of the IMC55 3’UTR region (Table S2). All primers utilized in this study are found in Table S2. To generate the plasmid pKD-IMC55-HA-apt, PCR products were transformed into the pKD *TetR* aptamer plasmid, digested with AatII and AscI, using NEBuilder HiFi DNA Assembly (New England Biolabs) <u>(Rajaram et al., 2020)</u>. All constructs used in this study had integration confirmed via Sanger sequencing. The IMC55^apt^ gRNA (oligo P7) was transformed into the pAIO #487 plasmid (pAIO-487-IMC55-apt) (a generous gift from the lab of Dr. Josh Beck, Iowa State University), digested with HindIII and AfLII, using the NEBuilder HiFi DNA Assembly (New England Biolabs) <u>(Glushakova et al., 2017)</u>.

The entire coding region of the IMC55 gene was recodonized (GeneWiz) and amplified using primers P10 & P11. Approximately ∼500 bp of 5’UTR homology region was amplified by primer pairs P12 & P13, and ∼500 bp of 3’UTR homology region was amplified by primer set P14 & P15, from NF54::DiCre genomic DNA. To generate the plasmid pH38-IMC55-loxPint-HA, a modified version of the pH38:pfikk10.1:loxPint:HA plasmid was first digested with EcoRI before transformation of the 5’UTR homology region and recodonized gene <u>(Jones et al. 2016)</u>. Next, the 3’UTR homology region was transformed into the plasmid, after digestion with AfeI <u>(Jones et al. 2016)</u>. The IMC55^KO^ gRNA (oligo P16) was transformed into the pAIO #487 plasmid (pAIO-487-IMC55-KO), digested with HindIII and AfLII <u>(Glushakova et al., 2017)</u>.

### Parasite culture and transfections

All parasite lines were maintained in a 1640 RPMI supplemented with AlbuMAX1 (Gibco) and incubated at 37°C with 5% CO_2_. Parasite cultures were kept at 2% hematocrit and transfected in duplicate as done previously <u>(Kudyba et al., 2018)</u>. For generation of IMC55^apt^ parasites, 25 μg of pKD-IMC55-HA-apt, containing the Blasticidin-S (BSD) drug cassette, and 25 μg of pAIO-487-IMC55-apt were transfected into NF54^attB^ parasites. Prior to transfection, pKD-IMC55-HA-apt was linearized overnight with EcoRV at 37°C. Transfected parasites were grown in 500 nM anhydrous tetracycline (aTc) (Cayman Chemical) for continuous expression of IMC55^apt^. To select for pKD-IMC55-HA-apt expression, drug pressure was applied 48 hours after transfection, using 2.5 μg/ml BSD. Once a mixed population had grown back, integration was confirmed using primer sets P1 & P5 and P1 & P6. Two independent clonal populations were generated using limiting dilution. Clonal IMC55^apt^ cultures were maintained in medium supplemented with 500 nM aTc and 2.5 μg/ml BSD.

To generate IMC55^KO^ parasites, 25 μg of pH38-IMC55-HA-DiCre, which contains the hDHFR resistance cassette and 25 μg of pAIO-487-IMC55-KO were transfected into NF54::DiCre parental line that expresses both halves of the rapamycin-dimerizing Cre recombinase <u>(Jones et al. 2016; Tibúrcio et al. 2019)</u>. Transfections were cultured in media supplemented with 0.5% AlbuMAX1 (Gibco). To select for pH38-IMC55-HA-DiCre, drug pressure was applied 48 hours after transfection using 2.5 nM WR99210

<u>(Collins et al. 2013; Jones et al. 2016)</u>. Drug selection was administered using a cycling method where parasites were cultured with WR99210 for four days, then cultured without drug pressure for four days until parasites grew back. Integration was validated using primer sets P8 & P9 and P8 & P15. Two independent clonal populations were generated using limiting dilution. Clonal IMC55^KO^ was maintained in 1640 RPMI supplemented with 0.5% AlbuMAX1 (Gibco).

### Synchronization assays

Parasite cultures were synchronized using a combination of sorbitol lysis and Percoll gradient (Genesee Scientific). Schizont-stage parasites were enriched using a 60% Percoll density gradient, spun at 2000 rpm for 15 minutes with no breaking. Percoll-enriched cultures were allowed to re-invade fresh new RBCs before isolation of ring-stage parasites. To selectively enrich ring-stage parasites, cultures were incubated in pre-warmed 5% sorbitol for 15 minutes at 37°C, before vigorous shaking and four rounds of washing in complete RPMI. After synchronization assays, parasites were cultured normally at 2% hematocrit.

### Immunoprecipitation assay

For PfERC immunoprecipitation experiments, PfERC mutants were generated as previously described <u>(Fierro et al. 2020)</u>. To identify interactors, > 10^9^ parasites from PfERC-*M9* or wild-type parental 3D7 (control) late schizonts were enriched via Percoll density gradient (Genesee Scientific) as described before, then collected in Extraction Buffer (40 mM Tris-HCl, 150 mM KCl, 1 M EDTA plus 1X HALT and 0.5% NP40) on ice for 15 minutes. Parasites were lysed via sonication then centrifuged at 20,000 xg at 4°C for 15 minutes. Lysates were incubated with anti-HA conjugated magnetic beads (Pierce) for 2 hours rocking at 4°C. Protein Loading Buffer (Li-COR Biosciences) was added, before samples were boiled for 5 minutes, then run on SDS-PAGE. The IP sample was cut from the SDS-PAGE gel and sent for mass spectrometry analysis to the Fred Hutch Institute (Seattle, Washington).

### PfERC invasion assay

Synchronized late PfERC-*glmS* and PfERC-*M9* schizonts were Percoll-isolated as described before (Genesee Scientific), then treated with 7.5 mM GlcN for 48 hours, and incubated with 20 μM E-64 for eight hours in an incubator at 37°C (Glushakova et <u>al. 2009)</u>. Merozoites were mechanically released from host RBCs by passing through a 1.2 μm Acrodisc Syringe Filter (PALL). Parasites were centrifuged at 2000 xg for 5 minutes, then resuspended in 100 μL of complete RPMI medium. Isolated merozoites were added to a 1 mL culture at 2% hematocrit. Parasites were grown in a FluoroDish cell culture dish (World Precision Instruments), shaking in a gassed chamber at 37°C for 20-24 hours. The rate of invasion was quantified using the following equation: 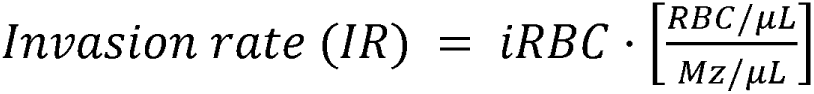. In this equation, “iRBC” is the final parasitemia at 20-24 hours, “RBC/μL” are the RBCs used before addition of merozoites, and “Mz/μL” are the free merozoites found in the 100 μL resuspended culture following syringe filtration. Each variable was measured via flow cytometry (CytoFLEX Beckman Coulter) and cells stained with Acridine Orange (ThermoFisher Scientific). To normalize data, the IR values for PfERC-*M9* were set to 100%. Flow cytometry data analysis was performed using FlowJo (Tree Star, Inc) and data were graphed using Prism (GraphPad Software Inc).

### Western blotting

For IMC55^apt^ western blot, synchronous parasites were washed six times in 1640 RPMI before being split into two conditions with one containing 500 nM aTc and 2.5 μg/ml BSD the other containing only 2.5 μg/ml BSD. For IMC55 samples, synchronous parasites were split into two conditions treated with either 100 nM rapamycin or 0.05% DMSO. Parasite pellets were collected and lysed every 24 hours with a 15 minute incubation in ice-cold 0.05% saponin in 1X PBS, followed by washing in ice-cold 1X PBS. Lysates were solubilized in protein loading dye supplemented with Beta-mercaptoethanol (Li-COR Biosciences) and loaded on SDS-PAGE.

Primary antibodies used for IMC55^apt^ were rat anti-HA (3F10; Roche; 1:2000) and rabbit anti-Ef1α (from Daniel Goldberg; 1:2000). Secondary antibodies used were IRDye 800CW goat anti-rabbit IgG and IRDye 800CW goat anti-rat IgG (Li-COR Biosciences; 1:20,000). Primary antibodies used for IMC55^KO^ were rabbit anti-HA (SG77; ThermoFisher Scientific; 1:400) and mouse anti-GAPDH (1.4; Millipore Sigma; 1:500). Secondary antibodies used were IRDye 680CW goat anti-rabbit IgG and IRDye 800CW goat anti-mouse IgG (Li-COR Biosciences; 1:20,000). Western blots were processed on the Odyssey Clx Li-COR infrared imaging system. Image analysis and quantification was performed in ImageStudio (Li-COR Biosciences).

### Microscopy and image analysis

Synchronized parasites were enriched for late schizonts via Percoll density gradient (Genesee Scientific) before collection for immunofluorescence assay (IFA). PKG-inhibited parasites were allowed to mature into late schizonts by addition of 1.5 μM Compound 1 (C1) {4-[2-(4-fluorophenyl)-5-(1-methylpiperidine-4-yl)-1H-pyrrol-3-yl] pyridine} for 4 hours before collection for IFA <u>(Collins et al. 2013)</u>. Parasites were smeared on a slide, then fixed with acetone for 10 minutes before washing three times in 1X PBS. Samples were blocked using 3% Bovine Serum Albumin (BSA) in 1X PBS. Antibodies used were rabbit anti-HA (SG77; ThermoFisher Scientific; 1:100), mouse anti-MSP1 (12.4; from Jana McBride via the European Malaria Reagent Repository; 1:500), rabbit anti-IMC1g (a gift from Jeff Dvorin; 1:2500), and rabbit anti-GAP45 (a gift from Jeff Dvorin; 1:1000). Secondary antibodies used were Alexa Fluor 488 and Alexa Fluor 546 (Life Technologies; 1:1000) in 3% BSA. Samples were mounted with Prolong Gold mounting media with DAPI (ThermoFisher Scientific) then imaged on the DeltaVision II microscope system with an Olympus Ix-71 inverted microscope. Images were taken as a Z-stack, deconvolved using SoftWorx, and displayed as a maximum intensity projection. Images were adjusted for display purposes only using Fiji.

### Growth assay

For the IMC55^apt^ growth assay, synchronous IMC55^apt^ ring-stage parasites were washed and split into plus and minus aTc conditions as described above. Cultures were placed in 96-well plates at 0.2% parasitemia at 2% hematocrit in triplicate and grown for 4 days. Samples were taken for growth assessment every 24 hours. At each timepoint, parasites were incubated shaking in 8 μM Hoechst (33342, ThermoFisher Scientific) for 20 minutes before parasitemia was assessed via flow cytometry on the Beckman Coulter CytoFLEX. Flow cytometry analysis was done using FlowJo (Tree Star, Inc) and data were graphed using Prism (GraphPad Software Inc).

For the IMC55^KO^ growth assay, synchronous ring-stage parasites were incubated with 100 nM rapamycin or 0.05% dimethyl sulfoxide (DMSO) for 16 hours before samples were taken every 48 hours to evaluate parasitemia. Cultures were grown in 96-well plate at 0.2% parasitemia and 2% hematocrit and grown for 7 days. Samples were incubated in Hoechst (33342, ThermoFisher Scientific) as described above and parasitemia was measured via flow cytometry on the Agilent Quanteon analyzer. Analysis was done as described above.

### Ultrastructural Expansion microscopy

Ultrastructural expansion microscopy (U-ExM) was done as before, with minor modifications <u>(Liffner and Absalon 2021; Anaguano et al. 2024)</u>. IMC55^KO^ late schizonts were enriched via Percoll density gradient (Genesee Scientific), then ML10-arrested (25 nM) (obtained from S. Osborne, BEI resources) for 3 hours as done before <u>(Ressurreição et al. 2020)</u>. Before collection, the culture was adjusted to 0.5% hematocrit, then incubated on a poly-D-lysine-treated 12 mm coverslip for 30 minutes at 37°C. Parasites were fixed with 4% paraformaldehyde in 1X PBS for 20 minutes at 37°C. A 1.4% paraformaldehyde/2% acrylamide (FA/AA) solution was added and incubated overnight at 37°C. After incubation, coverslips were washed three times in 1X PBS then placed cell-side up on parafilm in a humidity chamber on ice. The monomer solution (19% sodium acrylate, 10% acrylamide, 0.1% N,N’-methylenebisacrylamide in 1X PBS) was made within 10 days before use and stored at -20°C overnight before gelation. On ice, 5 μl of 10% ammonium persulfate (APS) and 5 μl of 10% tetramethylenediamine (TEMED) were added to 90 μl monomer solution and vortexed briefly. Quickly, 35 μl of the mixture were pipetted onto the parafilm and coverslip was placed cell-side facing down on the solution. Gels were incubated on ice for 5 minutes, then at 37°C for 30 minutes. Gels were transferred to a six-well plate with 2 mL of denaturation buffer (200 mM sodium dodecyl sulfate (SDS), 200 mM NaCl, 50 mM Tris, pH 9) and incubated rocking for 15 minutes at room temperature. Gels were then separated from the coverslip and transferred into a 1.5 mL tube filled with denaturation buffer, and incubated at 95°C for 90 minutes. Following denaturation, the gels were moved to a Petri dish with 25 mL of MilliQ H_2_O and incubated at room temperature for 30 minutes while rocking. The MilliQ H_2_O was replaced every 30 minutes for a total of three, 30 minute washes. The diameter of the gels were measured, then the gels were shrunk down with two 15 minute washes with 25 mL 1X PBS. Gels were transferred into a 24-well plate and blocked for 30 minutes with 3% BSA. Primary antibodies in 3% BSA were incubated overnight while shaking at room temperature, then washed three times in 0.5% PBS-Tween20 for 10 minutes. Secondary antibodies and specific stains were diluted in 1X PBS and incubated at room temperature while rocking for 2.5 hours. Gels were washed three times in 0.5% PBS-Tween20 for 10 minutes. Following staining, gels were expanded for a second time in a Petri dish with 25 mL of MilliQ H_2_O and incubated at room temperature for three, 30 minute washes while rocking. After expansion, gels were imaged immediately using a poly-D-lysine-treated glass-bottom imaging dish (Cellvis), or were stored in 0.2% propyl gallate at 4°C for future imaging.

Primary antibodies used were rat anti-HA (3F10; Roche; 1:50), rabbit anti-IMC1g (a gift from Jeff Dvorin; 1:250), and rabbit anti-GAP45 (a gift from Jeff Dvorin; 1:100). Secondary antibodies and stains used were Alexa Fluor 488 (Life Technologies; 1:500), Alexa Fluor 546 (Life Technologies; 1:500), SYTOX 647 (Invitrogen; 1:1000), NHS-Ester 405 (ThermoFisher Scientific; 1:250). Gels were imaged as Z-stacks on the Zeiss LSM 980 with Airyscan 2. Images were processed by the Airyscan using the Zen Blue software (Version 3.6). Representative images shown are maximum intensity projections of selected Z slices, with brightness and contrast adjusted for display purposes only.

## Acknowledgements

We thank Jeff Dvorin at the Boston Children’s Hospital at Harvard Medical School for anti-GAP45 and anti-IMC1g; Josh Beck at Iowa State University for the pAIO #487 plasmid; Moritz Treeck at the Gulbenkian Institute for Molecular Medicine for the pH38:pfikk10.1:loxPint:HA plasmid and the NF54::DiCre parasite line; Julie Nelson and Juan Bustamente at the CTEGD Cytometry Shared Resource Laboratory at the University of Georgia for help with flow cytometry and analysis; Muthugapatti Kandasamy at the Biomedical Microscopy Core at the University of Georgia for help with microscopy and image analysis; Fred Hutch Institute in Seattle, Washington for mass spectrometry analysis. This work was supported by grants from the U.S. National Institutes of Health to V.M. (R21AI133322 and R56AI173133) and M.A.F. and G.W.V. (T32AI060546).

## Supplemental Data

**Supplemental table 1.** Raw data from PfERC and control IP.

**Supplemental table 2.**
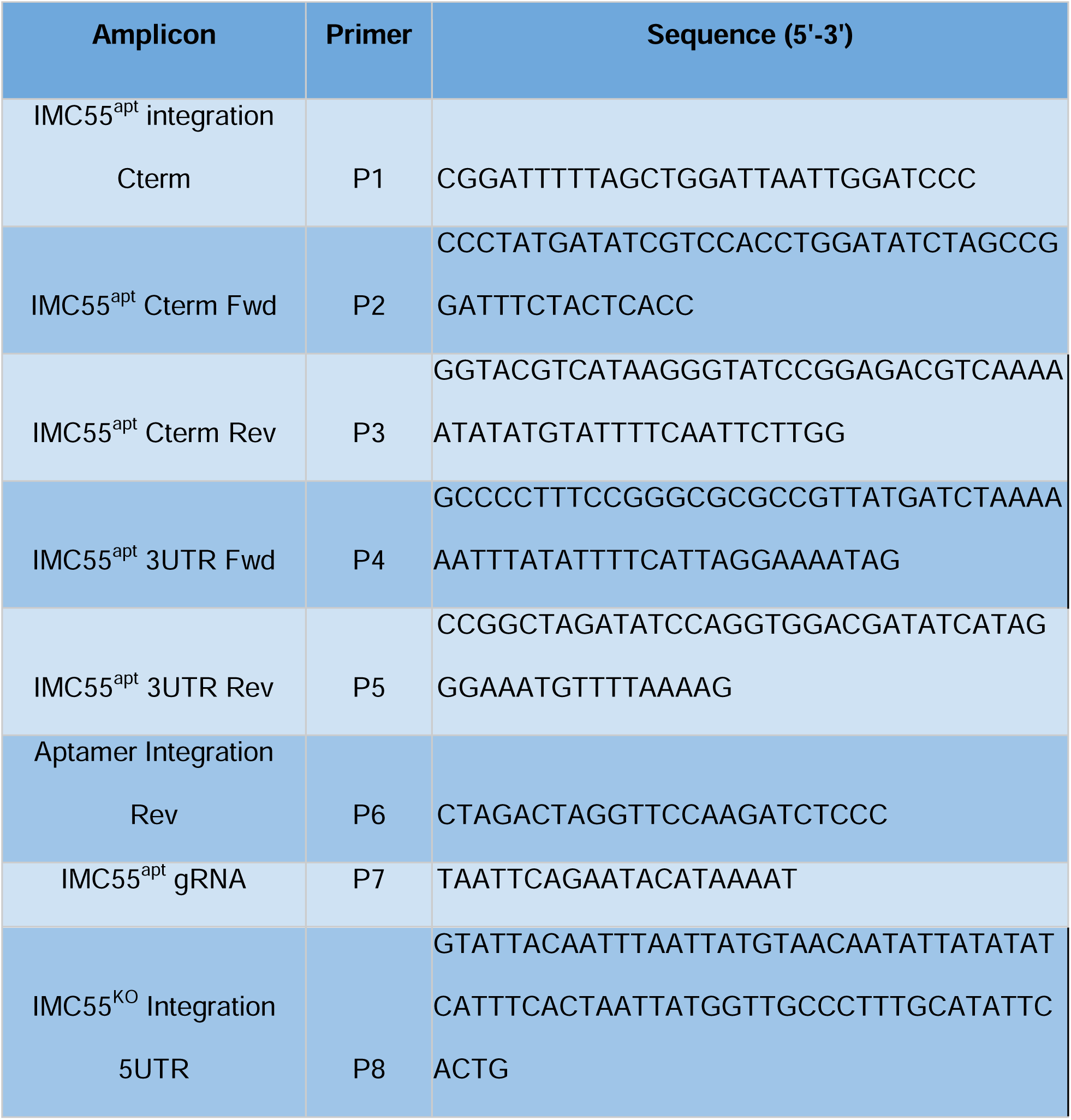
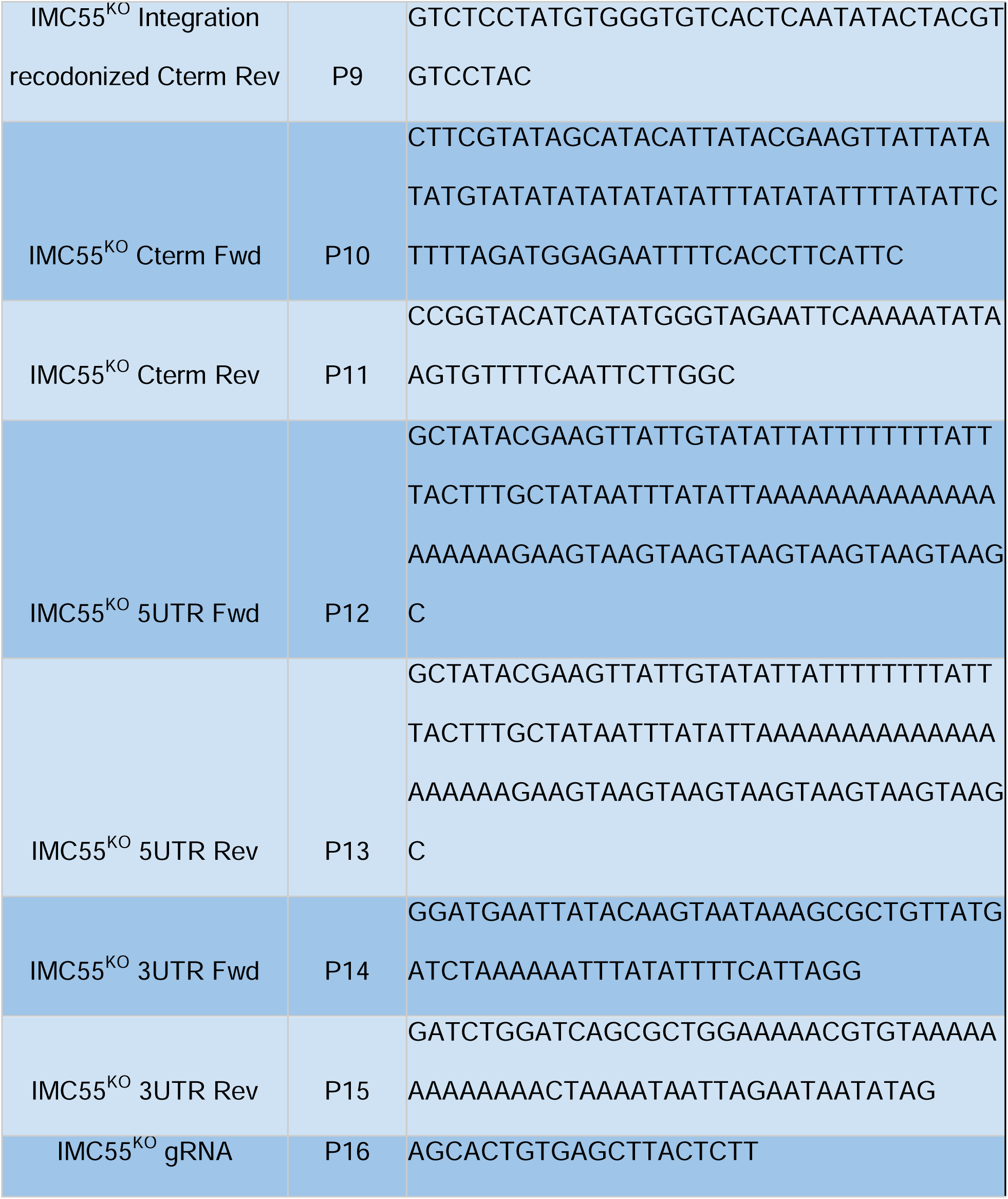
List of primers used to generate IMC55 conditional parasites in this study.

**Supplemental Figure 1.**
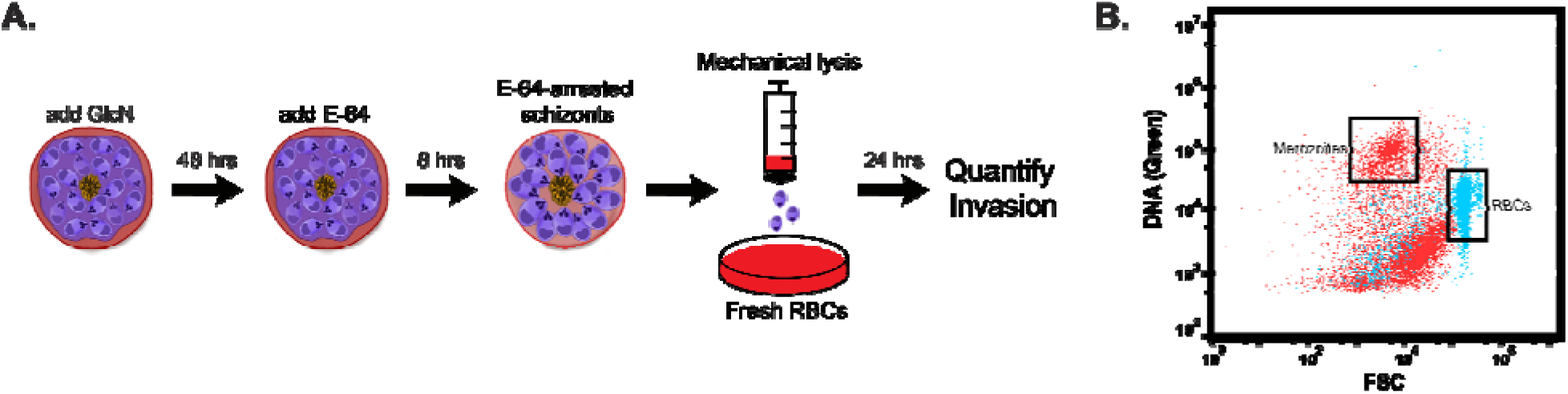
PfERC invasion assay. (A) Schematic showing isolation of infectious merozoites from RBCs and invasion assay. PfERC knockdown was E-64 treated for 8 hours, then parasites were mechanically lysed from host cells, and allowed to invade fresh RBCs. (B) Representative flow cytometry plot showing gating strategy to quantify merozoite (red) and RBC (blue) populations.

**Supplemental Figure 2.**
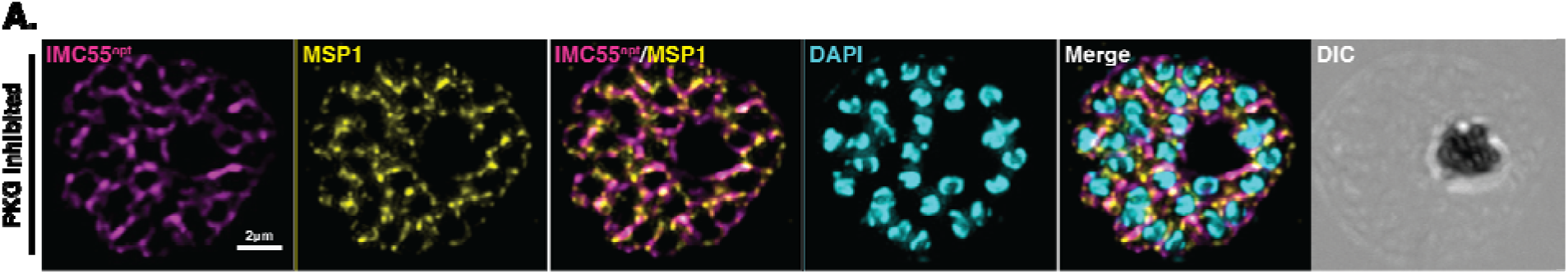
IMC55 localization under PKG inhibition. **(A)** Representative images from immunofluorescence assay showing subcellular localization of IMC55^apt^ late schizonts stalled using the PKG inhibitor C1 for 4 hours, then fixed via acetone and stained with specific antibodies. Antibodies used were anti-HA to detect IMC55^apt^ (magenta), a marker for the merozoite surface MSP1 (yellow), and the nuclear stain DAPI (cyan). Images include merged fluorescence and DIC. Images were acquired as a Z-stack, deconvolved, then projected as a single image. Representative images from 2 biological replicates.

**Supplemental Figure 3.**
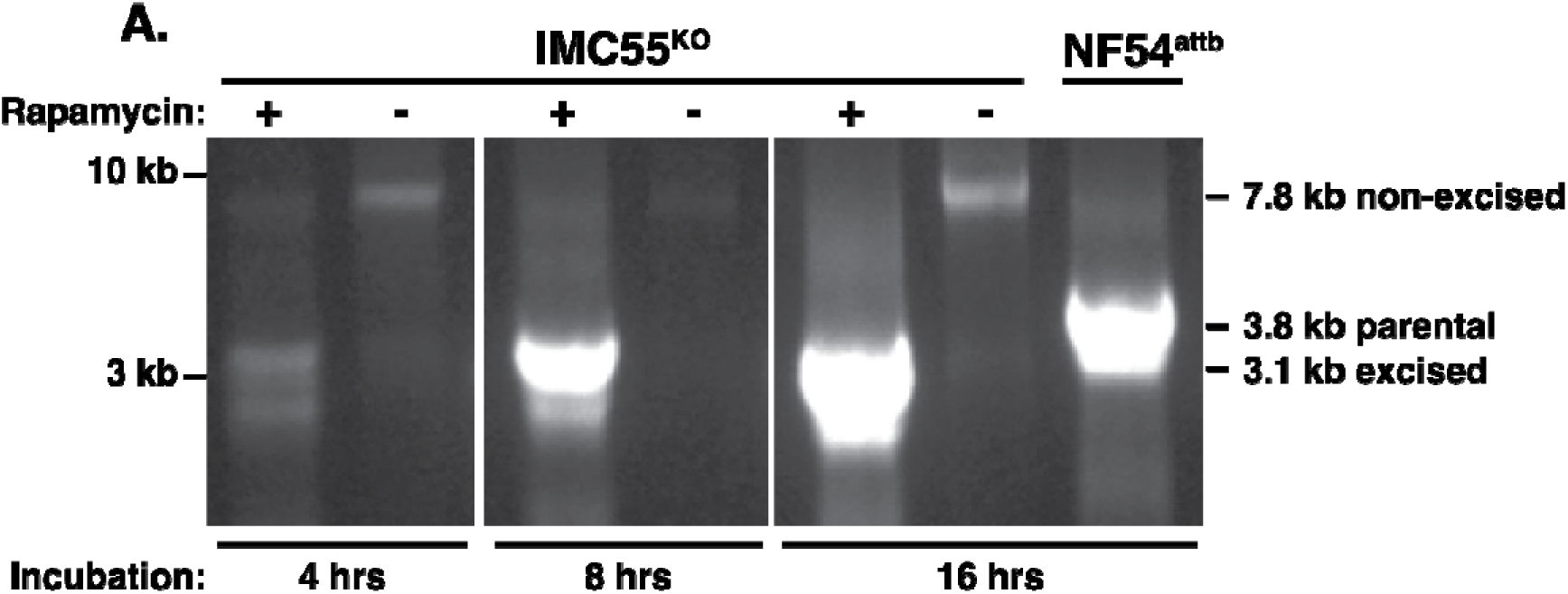
Optimization of rapamycin-mediated excision of IMC55^KO^. (A) PCR verification of IMC55^KO^ excision after incubation with 100 nM rapamycin or 0.05% DMSO for 4, 8, or 16 hours, collected immediately after RAP treatment. Amplification with primer pairs P8 & P15 generates a rapamycin-induced floxed IMC55^KO^ amplicon of 3.1kb, while treatment with DMSO produces a 7.8kb band amplicon. Parental NF54::DiCre amplification generates a 3.8kb amplicon. Representative data from 3 biological replicates.

**Supplemental Figure 4.**
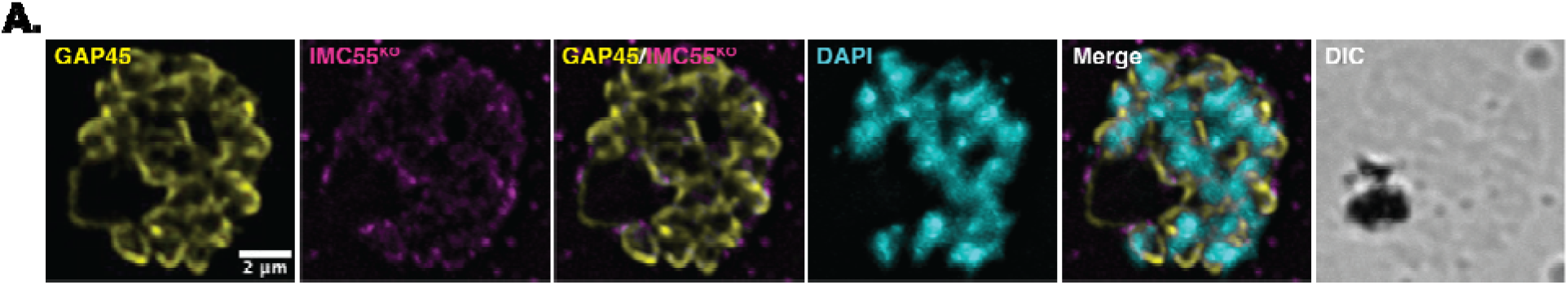
C**o**localization **of IMC55^KO^ with IMC markers.** (A) Representative images from immunofluorescence assay of IMC55^KO^ late schizonts fixed with acetone, then stained with specific antibodies. Antibodies used were anti-HA (magenta), anti-GAP45 (in yellow, top), and the nuclear stain DAPI (cyan). Images were collected as a Z-stack, deconvolved, then projected as a single image. Representative images from 3 biological replicates.

